# Chromosome-level *de novo* genome unveils the evolution of *Gleditsia sinensis* and thorns development

**DOI:** 10.1101/2024.01.11.575261

**Authors:** Dandan Xiao, Jiahao Liu, Jing Wang, Yuzhang Yang, Xiaoqian Yang, Ruen Yu, Chun Wang, Yanwei Wang, Yanping Liu, Dingchen Fan, Furong Lin

## Abstract

*Gleditsia* Linn is widely distributed in the world and *Gleditsia sinensis* as an important Leguminosae specie, especially its thorns, has been used in the Chinese medicine as a chief ingredient for centuries. While the lack of genome database limits its in-depth research, especially genetic regulation and molecular developmental mechanism. In this investigation, a chromosome-level *de novo* genome of *G. sinensis* was assembled through PacBio HiFi, Illumina sequencing as well as those from Hi-C, genetic mapping and K-mer analysis. The *G. sinensis* harbors 786.13 Mb sized genome (contig N50=1.58 Mb, scaffold N50=51.53 Mb, 2n=28) with 36, 408 protein-coding genes. The full-length transcriptome sequencing of diverse tissues was performed to assist genome functional annotation. The comparative and evolutionary analysis unveiled that *G. sinensis* diverged from the Cretaceous period approximately 76.31 million years ago (Mya) and the close relationship between *G. sinensis* and other 8 Leguminosae species. The whole-genome duplication (WGD) analysis indicated *G. sinensis* underwent three WGD events and might go through another WGD event after differentiating from other Leguminosae plants. The Weighted Gene Coexpression Network Analysis based on phenotype and differentially expressed genes further demonstrated that *GsinMYB* should be involved in the development of thorns via regulating late thorn differentiation. This investigation provides a high level genome of *Gleditsia* for Leguminosae species evolution comparison and functional elucidation and also key insights for further study on the molecular regulation mechanisms of the thorns development as a special abnormal stem organ in plants and the molecular breeding of *G. sinensis*.

## Introduction

*Gleditsia* Linn is widely distributed in the world, and *Gleditsia sinensis* Lam. (*G. sinensis*, 2n = 2x = 28) is a perennial deciduous tree belonging to Leguminosae sp. *G. sinensis* has been widely used in the traditional Chinese medicine as a chief ingredient for centuries (Lai et al., 2011; Zhang et al., 2016). Furthermore, the most medicinally active components of *G. sinensis* are mainly extracted from thorns and pods (Ji-Hye et al., 2018). The extract of thorns from *G. sinensis* has been reported to possess anti-angiogenic, anti-tumor, anti-cancer, antiseptic, and anti-inflammatory effects (Lee et al., 2010; Kai et al., 2016; Sujin et al., 2016; Zhang et al., 2016; Cai et al., 2019). While investigations involved in *G. sinensis* at the molecular level mostly focus on the pods. A miRNA expression profile of pods provided new insights into the molecular regulation of saponin synthesis in *G. sinensis* (Yang et al., 2022). The complete mitochondrial genome and chloroplast genome analysis provided the genetic evolution of *G. sinensis* and its phylogenetic relationship with other Fabaceae family members (Wei et al., 2020; Yang et al., 2021). However, to date, there were few investigations about the molecular mechanism of *G. sinensis* thorns.

As a special abnormal plant organ, multiple molecular regulation are involved in the development of the thorns. Genes expression profiling found that, *PIN7* and *LAX3* might be related with thorn phenotype of pummelo plant (Wu et al., 2021). Additionally, transcription factors such as TCP and NAC might regulate plant thorns development (Zhang et al., 2021; Zhang et al., 2021; Liu et al., 2023). With the increasing attention to the significant medicinal value of the *Gleditsia*, the molecular mechanism and gene regulation related to thorns development needs to be further elucidated. However, so far, little is known about the molecular regulation and the genes functions in *G. sinensis.* Despite of world-wide distribution and important medical and economic value of *G. sinensis*, there are no reports about its whole-genome sequences, which severely limited the in-depth exploration about the genetic regulation mechanisms related to medicinal components of thorns in trees with fast growth.

It is well-known that the first complete genome sequence of forest trees was reported in 2006 using *P*. *trichocarpa* (Tuskan et al., 2006). However, there are numerous repetitive sequences in forest tree genome, for example the genome of Chinese pine (*P*. *tabuliformis*) (Niu et al., 2022). The short read length of second-generation sequencing cannot effectively identify repetitive sequences and obtain information on full-length transcripts, which seriously restricts the assembly of complete and high-quality forest tree genome. However, in recent years, due to to the development of third-generation sequencing and the application of Hi-C technology, more than 200 high-quality tree genomes have been reported. Considering the importance of *G. sinensis*, its chromosome-level genome were assembled and harbors 786.13 Mb sized genome (contig N50=1.58 Mb, scaffold N50=51.53 Mb, 2n=28) with 36,408 protein-coding genes, combining PacBio long-reads, Illumina short-reads, and Hi-C sequencing and the full-length transcriptome sequencing of diverse tissues. The comparative and evolutionary analysis unveiled that *G. sinensis* diverged from the Cretaceous period approximately 76.31 million years ago (Mya) and the close relationship between *G. sinensis* and other 8 Leguminosae species, which diverged from *P*. *trichocarpa* and *A*. *thaliana* around 101.5 and 108.24 Mya respectively. The whole-genome duplication (WGD) analysis indicated *G. sinensis* underwent three WGD events (WGDⅠ, WGDⅡ and WGDⅢ) and might go through another WGD event after differentiating from other Leguminosae plants. Based on the high-quality genome map of *G. sinensis*, a sequential gene co-expression regulation network was performed combined phenotypic traits with the gene expression profiles of thorns among seven typical developmental periods during a natural growing cycle. Intriguingly, we found six differentially expressed genes, including genes encoding MYB transcription factor, which might be involved in regulating the development of thorns in *G. sinensis*. Comprehensively, this study provides a *de novo* high chromosome-level genome of *Gleditsia* for Leguminosae species evolution comparison and functional elucidation as well as a basis for the further research on molecular mechanisms underlying the development thorns as a special abnormal stem organ and molecular improvement of *G. sinensis*.

## Results

### *G. sinensis* genome sequencing and assembly annotation

To construct the genome sequence of *G. sinensis*, the datasets from PacBio HiFi and Illumina sequencing, as well as those from Hi-C, genetic mapping and K-mer analysis were comprehensively integrated. The genome size of *G. sinensis* was estimated to be approximately 709.43 Mb by K-mer analysis, with a repeat rate of 51.84 % and heterozygosity rate of 0.94% (Supplemental Fig. S1; Table 1). This genome assembly comprised 1329 contigs covering 786.13 Mb, with a contig N50 of 1.58 Mb, approximately 110.80% of the predicted genome size, of which 707.77 Mb located on 14 chromosomes, accounting for 90.04% of the assembled genome sequence (Fig. 1A; Table 1). The length of the sequence that could determine the direction was 665.82 Mb, accounting for 94.07% of the total length of the sequences located on chromosomes (Supplemental Table S1). The Hi-C interaction matrix exhibited a strong intra-chromosomal interactive signal along the diagonal, indicating the high quality of genome assembly (Supplemental Fig. S2). The BUSCO analyses showed the completeness of the genome to be 97.15%, the majority of them being single-copy genes, indicating that the integrity of gene prediction is high (Supplemental Table S2; Table 1).

**Figure 1.**
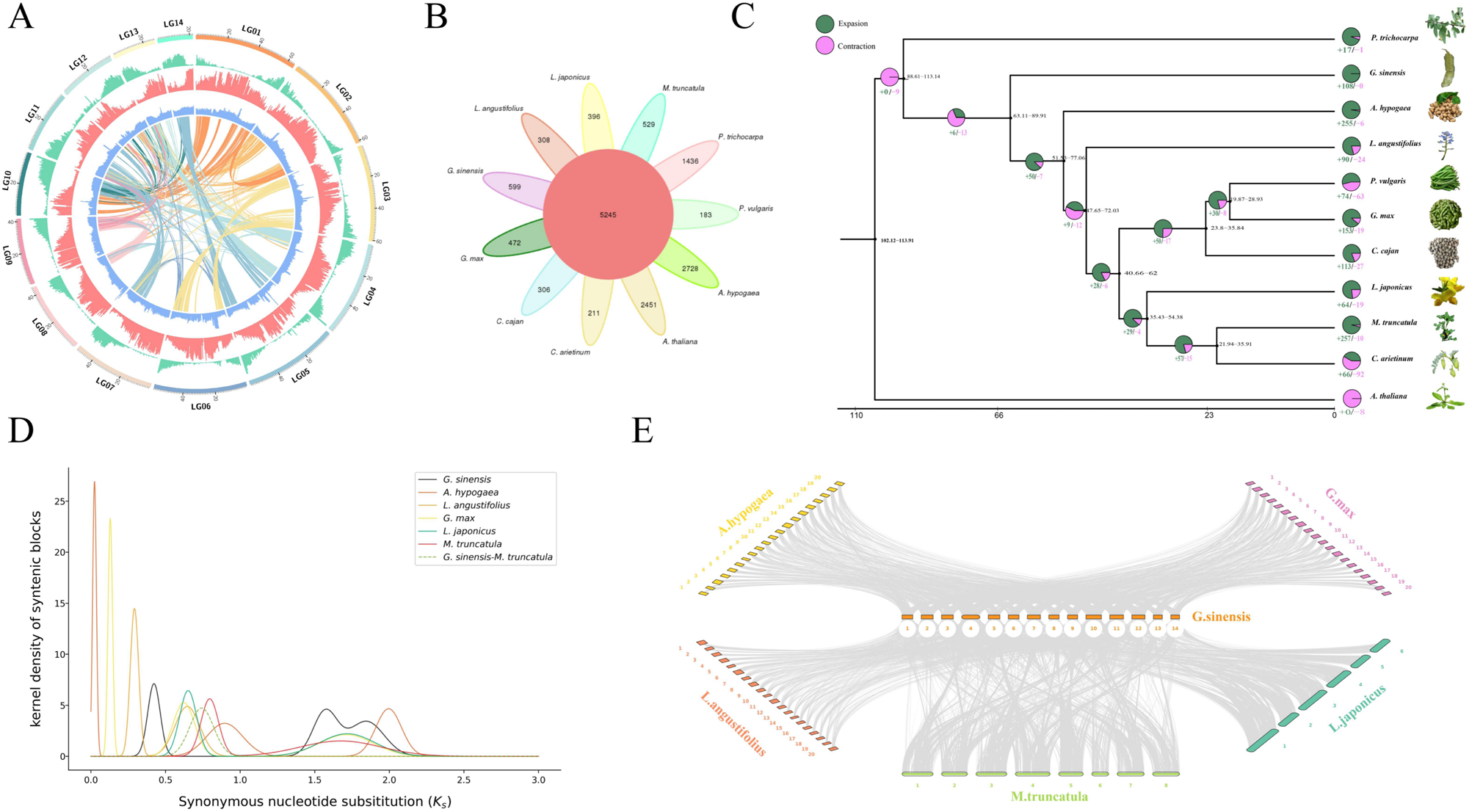
Thegenome characterization, phylogenomics and genome evolution of *Gleditsia sinensis*. **A)** Synteny and distribution of genomic features, **B)** Venn diagram exhibites the shared and unique gene families among *Cicer arietinum*, *Arachis hypogaea*, *Lotus japonicus*, *Phaseolus vulgaris*, *Glycine max*, *Cajanus cajan*, *Lupinus angustifolius* and *Medicago truncatula*, *Populus trichocarpa*, *Arabidopsis thaliana* and *Gleditsia sinensis*. **C)** The phylogenetic tree, divergence times and gene family expansion and contraction among 11 plant species. (‘+’ represents the number of gene families that expand on this node, and ‘−’ represents the number of gene families that contract on this node. The pie chart represents the proportion of corresponding branch contraction and expansion gene families. The numbers next to the node indicate the range of species divergence times. The bottom line represents the timeline, unit: million years ago). **D)** Ks distribution of *Populus trichocarpa*, *Gleditsia sinensis*, *Arachis hypogaea*, *Lupinus angustifolius*, *Phaseolus vulgaris*, *Glycine max*, *Cajanus cajan*, *Lotus japonicus*, *Medicago truncatula*, *Cicer arietinum*, and *Arabidopsis thaliana*. **E)** Chromosome collinearity analysis among *Arachis hypogaea*, *Lupinus angustifolius*, *Glycine max*, *Lotus japonicus*, *Medicago truncatula* and *Gleditsia sinensis*.

**Table 1.**
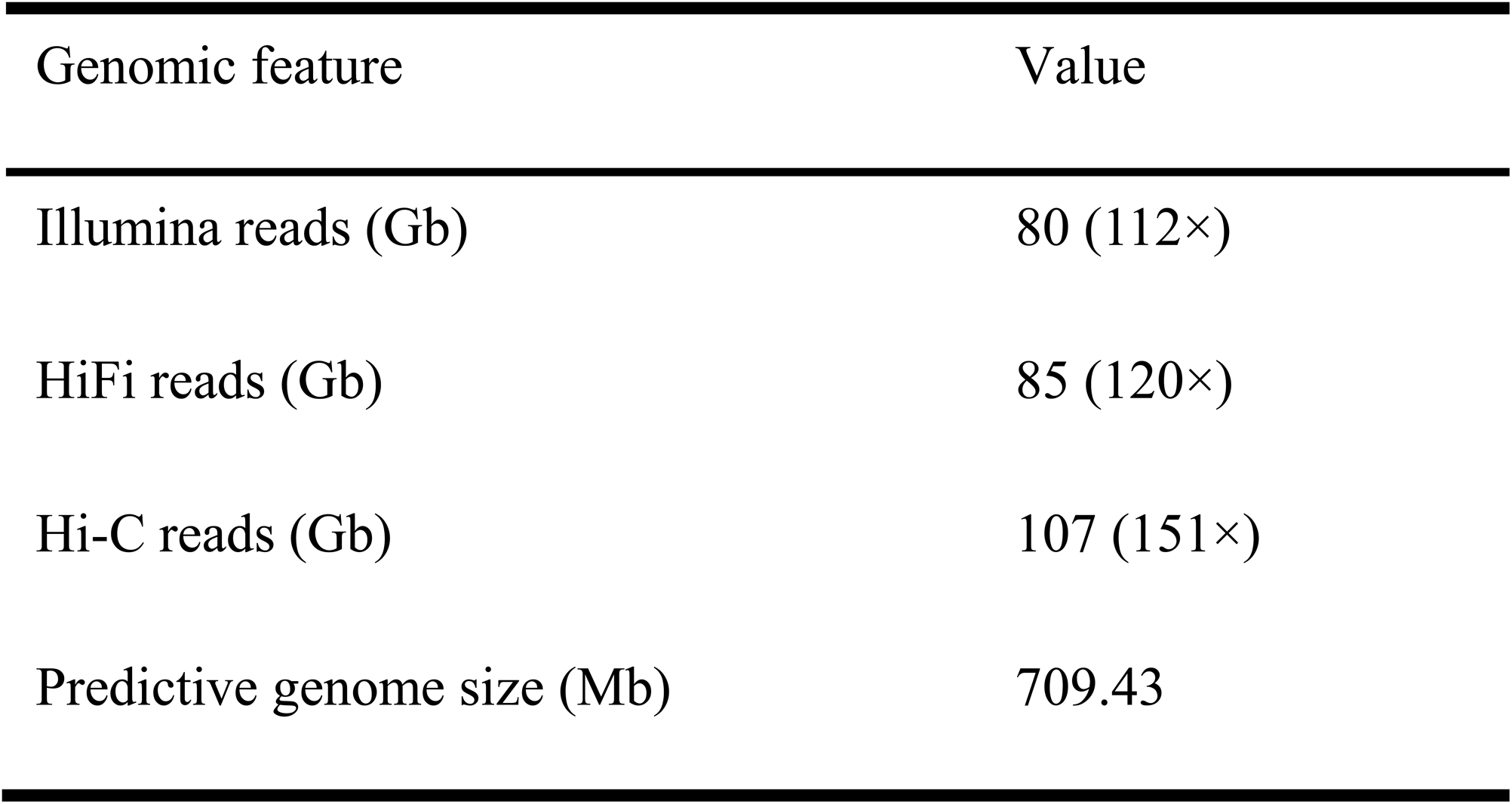
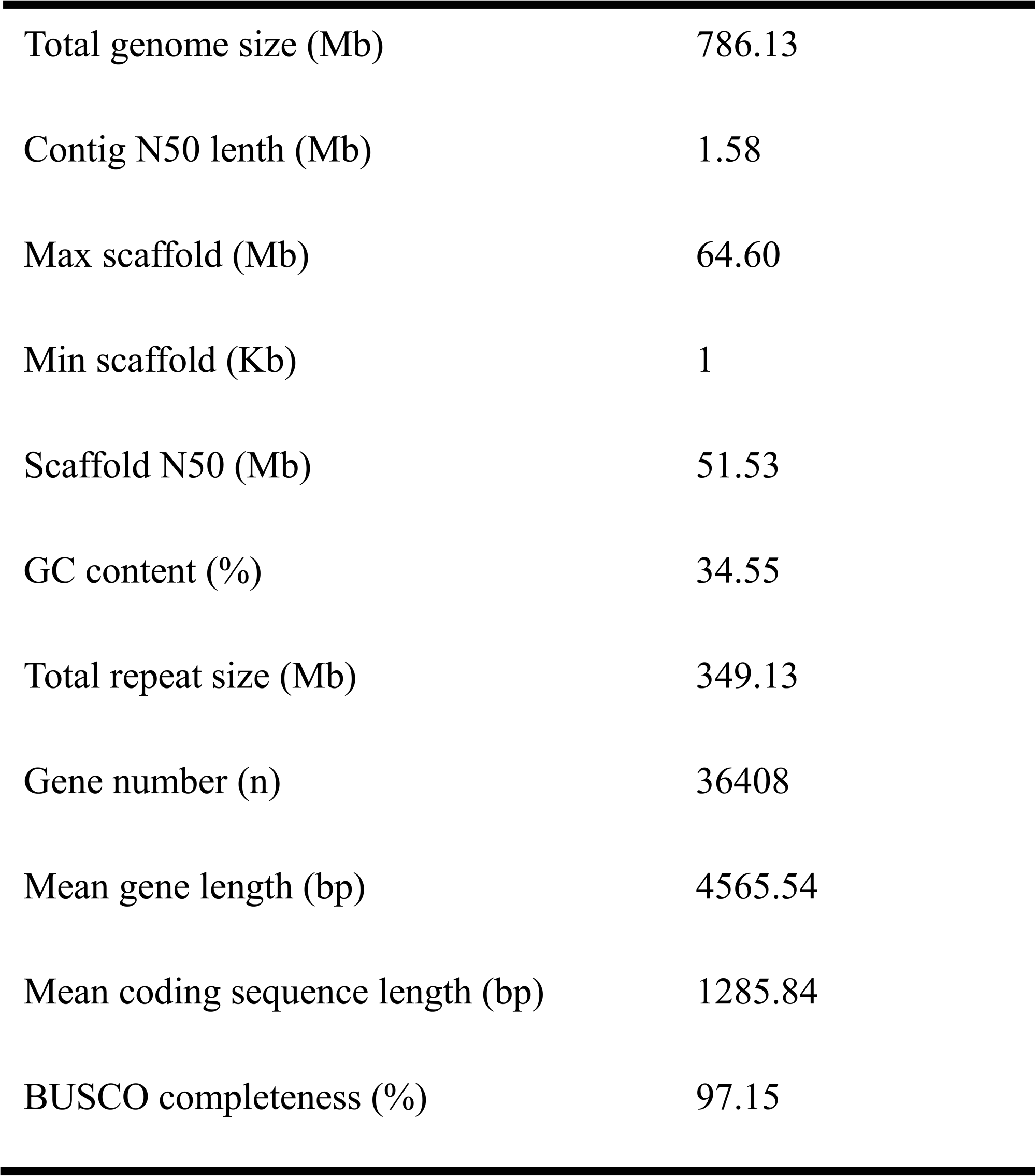
Statistics for the *Gleditsia sinensis* genome sequencing and assembly.

### *G. sinensis* genome annotation

A total of 349.13 Mb of repeats were identified, accounting for 44.41% of the assembled *G. sinensis* genome (Table 1). The dominant was transposable elements (TEs), accounting for 42.86% with 336.94 Mb in genome (Supplemental Table S3). Further analysis indicated that retroelement was significantly higher compared to DNA transposon. The dominant retroelement were LTRs (243.8 Mb, 31.02%) with Gypsy (18.34%), followed by unknown LTRs (6.76%) (Supplemental Table S3). Additionally, there were approximately 12.19 Mb tandem repeats, accounting for 1.55% of the genome (Supplemental Table S4).

The combination of *de novo*, homology-based prediction and transcriptome-based approaches was used to annotate the protein-coding genes. RNA-sequencing libraries were built for gene annotation. Finally, a total of 36408 genes were detected and predicted, and the average lengths for gene length, coding sequence (CDS), exon sequence, and intron sequence were 4565.54, 1285.84, 1597.61, 2967.93 bp, respectively (Supplemental Table S5). Additionally, 17,796 non-coding RNAs were identified, including 15,473 ribosomal RNAs (rRNA), 961 transfer RNAs (tRNA), 179 microRNA (miRNA), 587 small nuclear RNAs (snRNA), 596 small nucleolar RNA (snoRNA) (Supplemental Table S6). 248 pseudogenes were predicted, the total length and average length were 1,240,393 bp and 5001.58 bp, respectively (Supplemental Table S7). Among protein coding genes, 34,674 (95.24%) genes could be annotated by at least one of the following database: TrEMBL protein database (93.81), NCBI non-redundant protein database (NR) (93.11%), the SwissProt protein database (75.57%), Gene Ontology (GO) (76.45%), KEGG PATHWAY (70.92%), KOG (51.17%), EggNOG (79.86%), and the protein families database (Pfam) (80.67%) (Supplemental Table S8). In addition, there were 1,405 motifs and 36,170 domains were annotated.

### Comparative and evolutionary analysis of *G. sinensis*

To explore the phylogenetic relationship and evolutionary status of *G. sinensis* genome, the genome of *G. sinensis* was compared to 8 Leguminosae species (*C*. *arietinum*, *A*. *hypogaea*, *L*. *japonicus*, *P*. *vulgaris*, *G*. *max*, *C*. *cajan*, *L*. *angustifolius* and *M*. *truncatula*) and other 2 model plants (*P*. *trichocarpa* and *A*. *thaliana*) (Fig. 1B). A total of 32,785 *G. sinensis* genes were divided into 18,634 gene families, including 599 specific gene families of *G. sinensis* and 5,245 gene families shared by all species through gene family cluster analysis (Supplemental Table S9). Furthermore, the specific gene families of *G. sinensis* were mainly enriched in ‘Cytokinin biosynthetic process’ and ‘External encapsulating structure’ through GO analysis, and were mainly enriched in ‘Purine metabolism pathway’, ‘RNA transport pathway’ and ‘Galactose metabolism pathway’ through KEGG analysis, indicating these specific gene families may be involved in the growth and development, formation of thorns in *G. sinensis*.

The 1,238 single-copy gene families from these 11 species were used to reconstruct phylogenetic relationship. The maximum-likelihood (ML) tree showed that *G. sinensis* diverged from the Cretaceous period approximately 76.31 million years ago (Mya), indicating the close relationship between *G. sinensis* and 8 other Leguminosae species, including the closest relationship between *G. sinensis* and *A*. *hypogaea*. Additionally, the 8 Leguminosae species diverged from *P*. *trichocarpa* and *A*. *thaliana* around 101.5 and 108.24 Mya (Fig. 1C).

Comparing other 10 plant species, there were 108 gene families were expanded. More intriguingly, GO functional enrichment analysis found these expanded gene families of *G. sinensis* were significantly enriched in sesquiterpenoid and triterpenoid biosynthesis, and selenocompound metabolism (Supplemental Fig. S3A). Additionally, functional enrichment analysis revealed the expanded gene families were significantly enriched in sesquiterpenoid and triterpenoid biosynthesis pathway, and selenocopound metabolism (Supplemental Fig. S3B). To investigate the positively selected genes of *G. sinensis*, the single copy gene families among *L*. *japonicus*, *C*. *cajan*, *G*. *max*, *L*. *angustifolius* and *G. sinensis* were selected for positive selection analysis using PAML. As a result, 45 genes were positively selected, and their molecular function were enriched in fruit development, seed development, embryo development, embryo development ending in seed dormancy and DNA recombination, indicating their significant implications for the generation of new functions in *G. sinensis*.

### Whole-genome duplication in *G. sinensis*

The whole-genome duplication (WGD) events were unveiled through Ks and 4DTv analyses. The density of Ks for *G. sinensis* presented three peaks in Ks≈0.42 and Ks≈1.5-2.0 respectively, which indicated *G. sinensis* underwent three WGD events (WGDI, WGDII and WGDIII). Moreover, the time points of WGDI, WGDII in *G. sinensis* were similar to the WGDI of *A*. *hypogaea*, *G*. *max*, *L*. *japonicus* and *M*. *truncatula*, which suggested all these Leguminosae species underwent a WGD event. Additionally, the two peaks in Ks≈1.5-2.0 were similar and continuous, which suggesting the WGDI and WGDII might represent a WGD event. Intriguingly, the Ks peak of *G. sinensis-M*. *truncatula* was located before the Ks peak of WGDIII in *G. sinensis*, implying *G. sinensis* underwent another WGD event after differentiating from other Leguminosae plants (Fig. 1D).

The intragenomic chromosome collinearity analysis identified 814 collinear gene blocks in *G. sinensis*, including 12893 collinear genes (Supplemental Table S10), proving the occurrence of polyploidization events in the *G. sinensis* genome. Additionally, the comprehensive analysis of chromosome collinearity analysis and Ks values caculation of one-vs-one orthologs of *G. sinensis*-*A*. *hypogaea*, *G. sinensis*-*L*. *angustifolius*, *G. sinensis*-*G*. *max*, *G. sinensis*-*L*. *japonicus*, *G. sinensis*-*M*. *truncatula* showed these Leguminosae species (*G. sinensis*, *L*. *angustifolius*, *G*. *max*, *L*. *japonicus* and *M*. *truncatula*) have a close evolutionary relationship (Fig. 1E; Supplemental Fig. S4).

### Phenotype characters of thorns

To clearly elaborate the phenotype characters of thorns in *G. sinensis*, the development time were divided into 8 stages in this investigation using characters such as colors and shapes, including formative stage (GSB1, August to early November), dormancy stage (GSB2, early November to late March of the following year), germination stage (GSB3, late March), the stage of buds transforming to thorns (GSB4, the first ten-day period of April), fast-growing stage (GSB5, mid to late April), late growth stage (GSB6, May to mid July), browning stage (GSB7, mid July to August) and maturity stage (GSB8, September to December). There were brown scale coated buds appeared on stem protrusion at GSB1 (Fig. 2, A and B). However, no obvious changes of colors and shapes were observed at GSB2 at early January developed from GSB1 at late October (Fig. 2, C and D). While the buds began to sprout at GSB3 at early April (Fig. 2E) and quickly transformed into tender thorns, around which there were branches of thorns and leaves at GSB4 at mid-April (Fig. 2F). The length and size of thorns changed obviously at GSB5 at late April (Fig. 2, G and H). The growth rate of thorns slowed down or stopped growing in length GSB6 at mid-June (Fig. 2I). The thorns were becoming hard gradually at GSB7 at early August (Fig. 2J). The color of thorns changed from green to yellow brown and then to purple brown, furthermore, the thorns could persist on branches at GSB8 (Fig. 2H). To clarify the development characters of thorns in *G. sinensis*, the length, basal diameter and weight were calculated at GSB1-GSB7. The thorns developed slowly from GSB1 to GSB3, then appeared rapid growth from GSB4 until reached peaks at GSB7. Overall, there were significant differences in the length and weight but no significant change in basal diameter as the development stage changed (Table 2).

**Figure 2.**
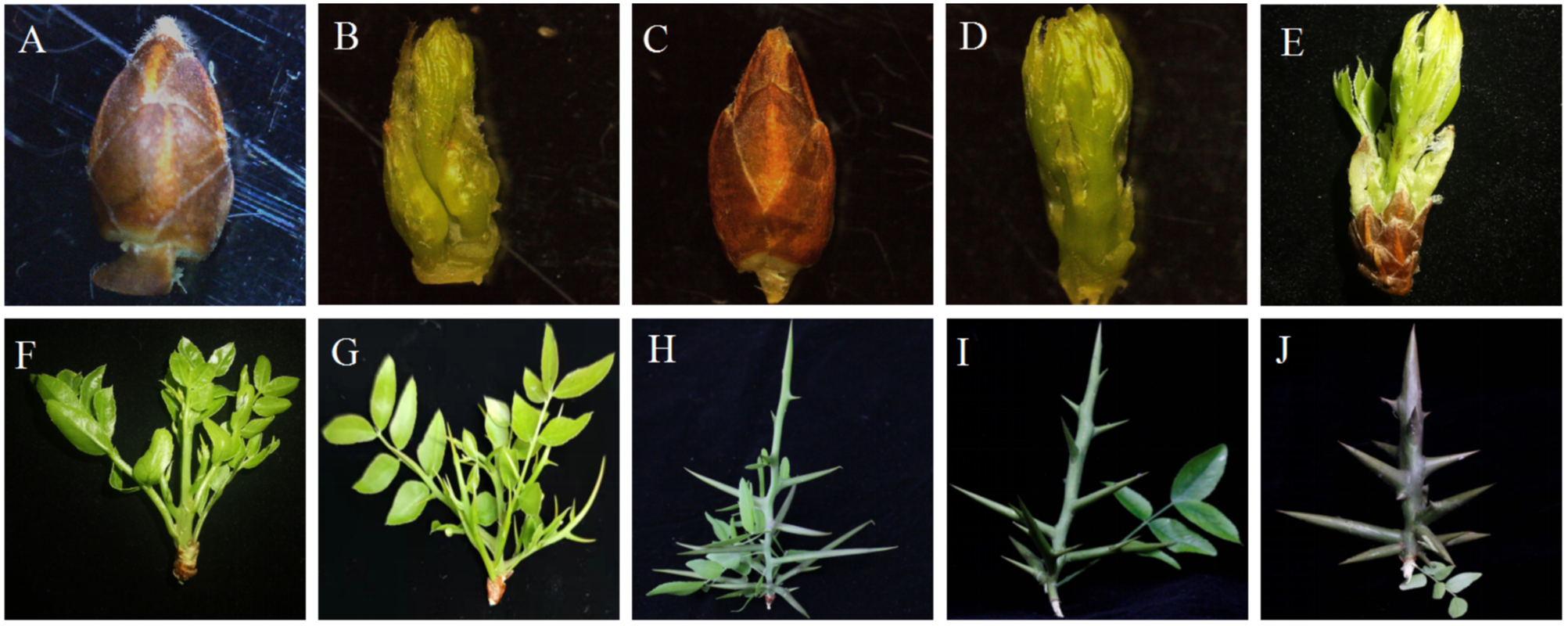
Phenotype of seven developmental stages of *Gleditsia sinensis* thorns. **A)** and **B)** the buds with or without the scale leaf coating at the formative stage GSB1, **C)** and **D)** the buds with or without the scale leaf coating at the dormancy stage GSB2, **E)** the buds at the germination stage GSB3, **F)** the buds at the stage of buds transforming to thorns GSB4, **G)** and **H)** the thorns at fast-growing stage GSB5 with or without nutrient leaf, **I)** the thorns at late growth stage GSB6, **J)** the thorns at the browning stage GSB7.

**Table 2.**
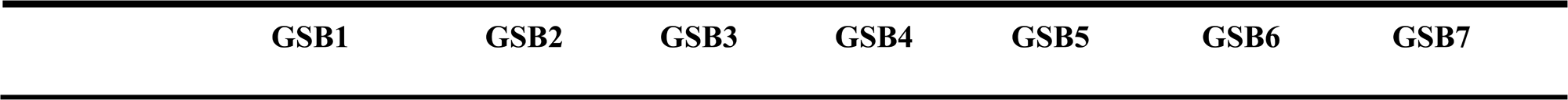
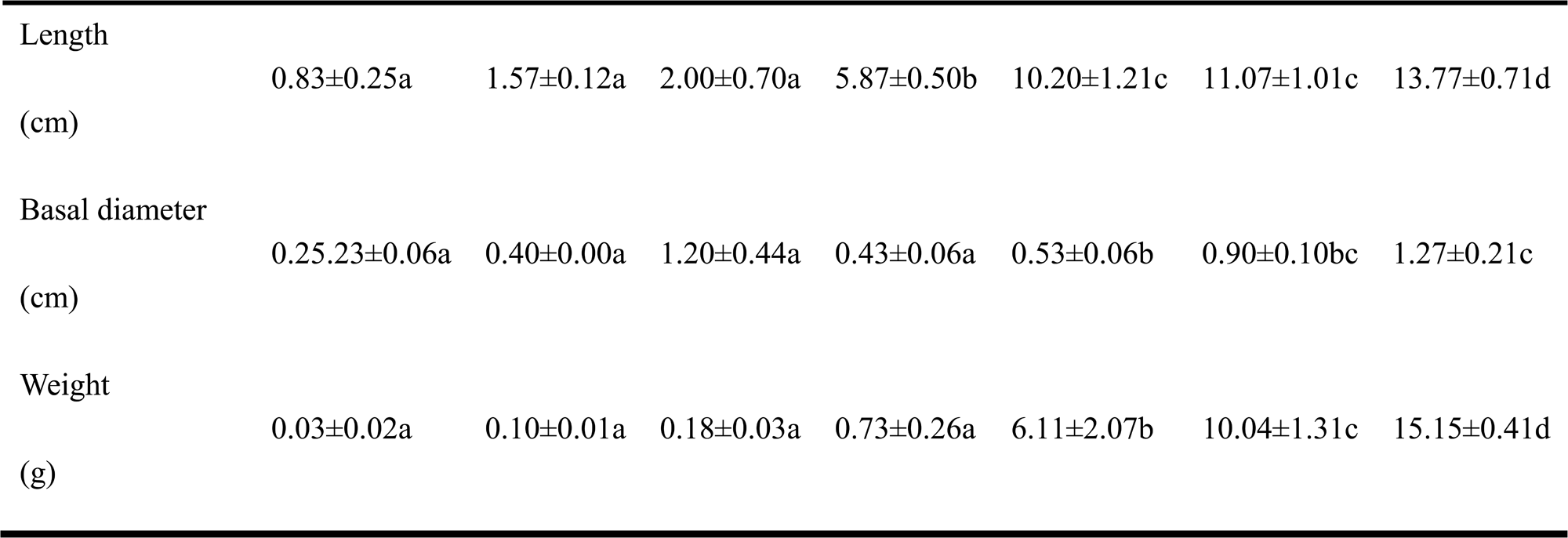
The length, basal diameter and weight of the seven development stages of *Gleditsia sinensis*.

### Anatomical characteristics of thorns

The detailed microstructure and developmental changes of thorns from GSB1 to GSB3 were further investigated to analyze the anatomical differences, including longitudinal sections and transverse sections of thorns. The SEM analysis revealed that the scale bud were tightly wrapped by multiple layers of scale leaves, and the outer layer of scale leaves had been embolized in stem protrusion. Additionally, there were many filamentous material around the scale leaves at GSB1 (Fig. 3, A and E). We observed more obvious squamosal leaf primordia and filamentous material at GSB2 (Fig. 3, B and F). At incipient germination stage GSB3, the spines broke through the scales and began to germinate, including the terminal bud and many lateral buds (Fig. 3, C and G). The elongation of terminal bud and lateral buds could be observed at late budding stage GSB3 (Fig. 3, D and H).

**Figure 3.**
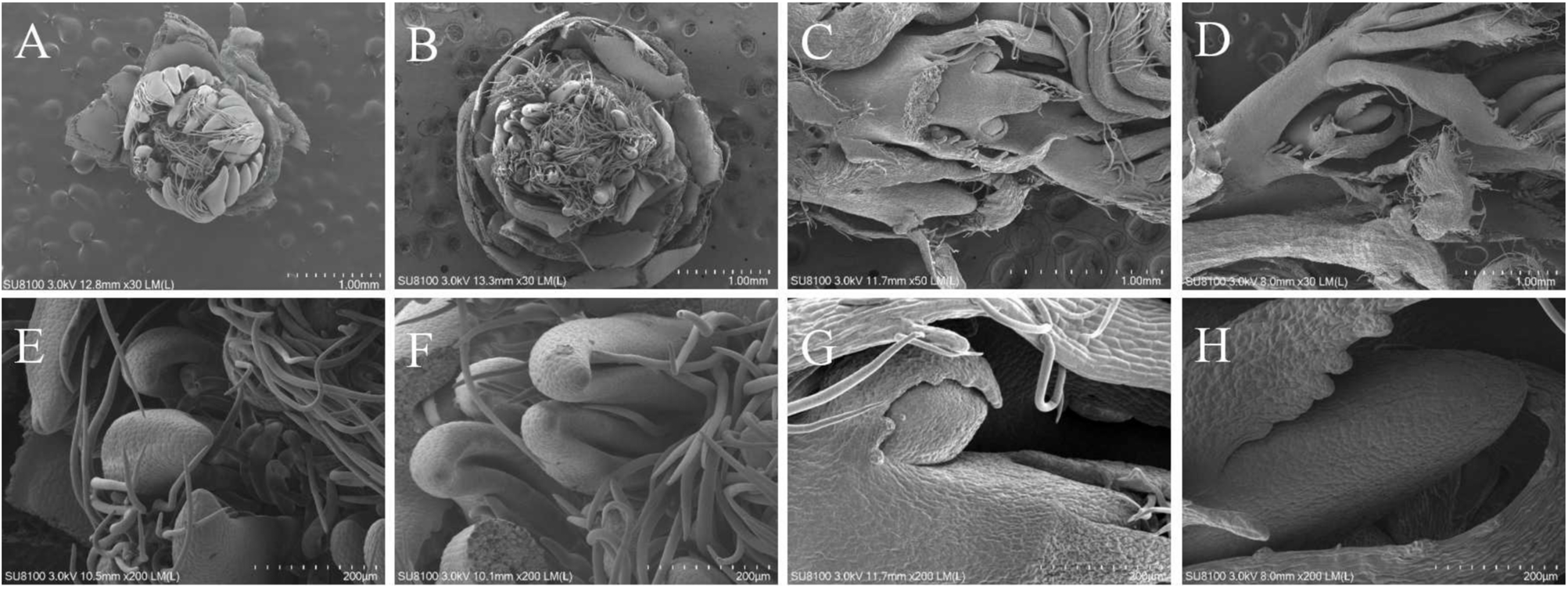
The scanning electron microscope observation at different developmental stages of *Gleditsia sinensis* thorns. **A)** and **E)** GSB1, **B)** and **F)** GSB2, **C)** and **G)** GSB3, **D)** and **H)** GSB4, scales of **A)**, **B)**, **C)**, and **D)** are 1 mm, scales of **E)**, **F)**, **G)**, and **H)** are 200 um.

The paraffinembedded sections observation revealed the basic structure of the bud had formed at GSB1 (Fig. 4, A and D). The growth cone, bud primordia, scale leaf, scale leaf primordia and axillary bud primordia can be observed at longitudinal section (Fig. 4D). The structure of epidermis, cortex, phloem, cambium, xylem and pith can be seen from the outside to the inside on the bud axis in the transverse section, and the structure of central area mother cell, central tissue center, costal meristem area and peripheral meristem area can be seen in the meristem area (Fig. 4A). However, there was no obvious development about meristems of stem thorns at GSB2, which was consistent with the phenotypic changes of thorns and could be regarded as a transition phase (Fig. 4, B and E). The thorns had formed a primary structure, including meristematic zone, elongation zone and mature zone at longitudinal section. Furthermore, the main structures of the mature area were epidermis, cortex, pericycle, primary phloem, cambium, primary xylem and pith from outside to inside. And the leaf primordia were observed in the axillary bud site (Fig. 4E). As growth proceeded in the thorn primordium, the various primordia and other parts began to grow and gradually transformed from buds to thorns at GSB3 (Fig. 4C and F). These results suggested the stem spines were evolved from the branch spines.

**Figure 4.**
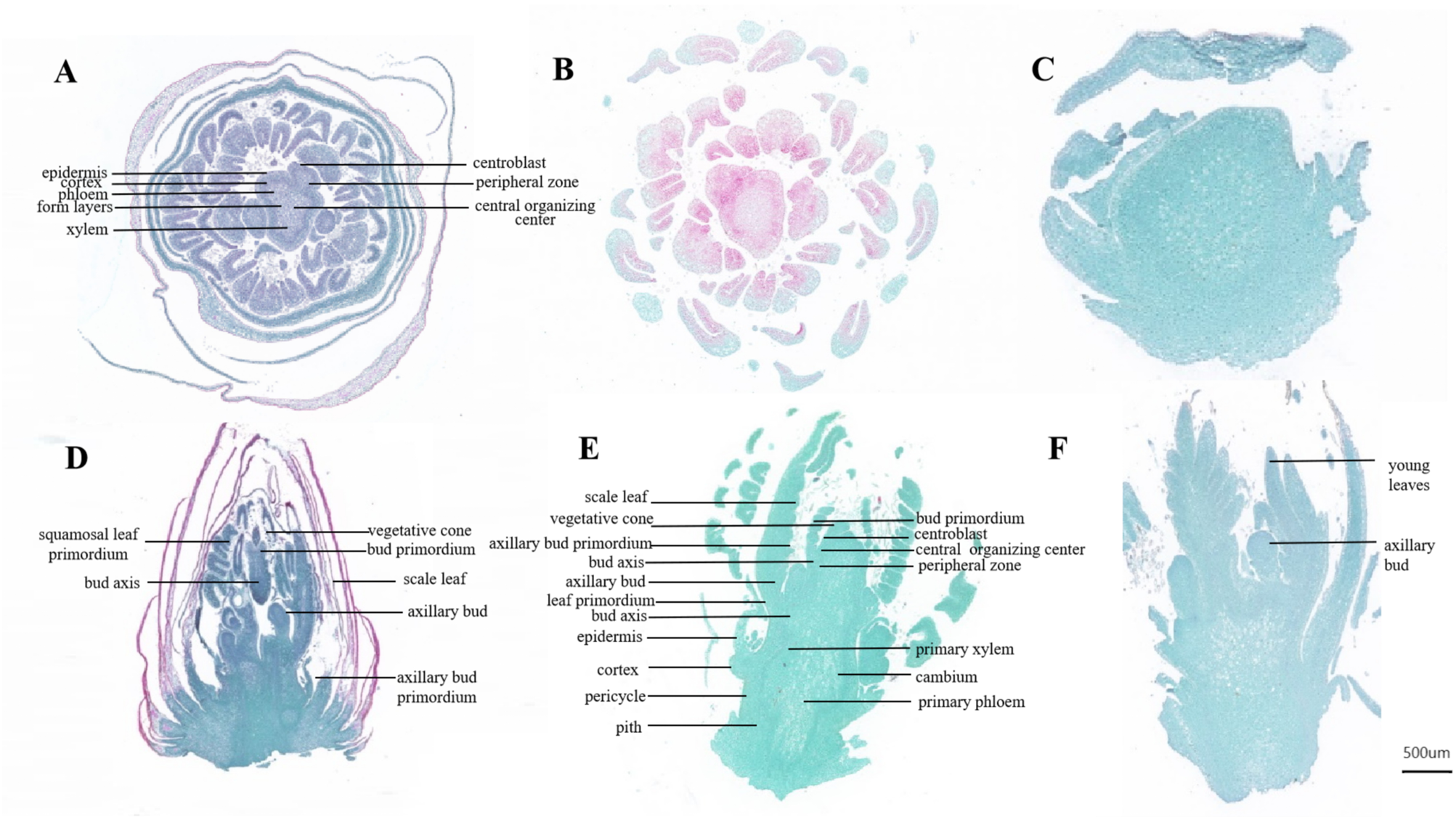
Observation on the microstructure of thorns of main stem at different developmental stages of *Gleditsia sinensis* thorns. **A)** and **D)** represent the transverse and longitudinal sections during GSB1, **B)** and **E)** represent the transverse and longitudinal sections during GSB2, and **C)** and **F)** represent the transverse and longitudinal sections during GSB3.

### Gene expression profiles during thorns development of *G. sinensis*

To characterize the molecular regulation of thorn development activity, the gene expression profiles of thorns at seven development stages were further examined. A total of 133.66 Gb clean data were yielded from 21 cDNA libraries. The Q20 and Q30 quality scores attained at least 98.12% and 94.24% respectively, and the overall GC content were more than 44.02% (Supplemental Table S11). The alignment efficiency of sequencing reads was between 92.74% and 95.23%, which further supported the high assembling completeness of *G. sinensis* genome. Based on FPKM (Supplemental Fig. S5A), statistical dispersion of gene expression level distribution (Supplemental Fig. S5B), pearson correlation r analysis (Supplemental Fig. S5C), and principal component analysis (PCA, Supplemental Fig. S5D), the standardized analysis proved that the accuracy of RNA-seq among thorns with different development stages and could be used for subsequent analysis.

### Investigation of DEGs during thorns development in *G. sinensis*

To investigate the genes involved in regulating the thorns development, we examined the DEGs between every two developmental stages. The comparative analysis showed that there were more DEGs (11,866 DEGs) between GSB3 and GSB2 among other comparison groups, followed by GSB5 vs GSB2 (11,269 DEGs) and GSB7 vs GSB3 (11190 DEGs) (Supplemental Fig. S6A). It indicated that there was a maximum number of DEGs expression regulation during the thorns transition from dormancy stage (GSB2) to germination stage (GSB3) compared with other two developmental stages. Interestingly, we found there was no common DEGs among every two developmental stages, indicating there was no DEGs regulating the whole developmental progress. Additionally, GSB7 vs GSB3 had the maximum number of specific DEGs (190 DEGs), followed by GSB3 vs GSB2 (155 DEGs), GSB4 vs GSB2 (145 DEGs), GSB5 vs GSB2 (144 DEGs) and GSB4 vs GSB1 (102 DEGs) (Supplemental Fig. S6B). The DEGs comparison between adjacent developmental stages supported that there were 15 common DEGs, GSB2 vs GSB1 had most specific DEGs (n=4,559), followed by GSB5 vs GSB4 (n=1,158) and GSB3 vs GSB2 (n=729) (Supplemental Fig. S6C). These results indicated that the thorns development was slow at GSB2 and GSB3 with relatively fewer DEGs compared to other stages, which was consistent with phenotypic changes of thorns. To explore the functions of DEGs involved in thorns development, GO annotation and KEGG enrichment investigation were performed in adjacent developmental stages. For biological process, the most highly enriched GO terms was ‘Carbohydrate metabolic process’ respectively in GSB2 vs GSB1 and GSB6 vs GSB5, suggesting the important role of carbohydrate metabolism in the thorns transition from formative stage to dormancy stage, and from fast-growing stage to late growth stage. In GSB3 vs GSB2 and GSB4 vs GSB3, the most highly enriched GO term was ‘Response to abiotic stimulus’, suggesting the abiotic stimulus affected the development of thorns from dormancy stage to the stage of buds transforming to thorns. The GO term ‘Oxidation-reduction process’ was most highly enriched in GSB5 vs GSB4, suggesting its important regulation role from the stage of buds transforming to thorns to fast-growing stage. Additionally, the maximum number of DEGs were enriched in GO term ‘Response to stimulus’ in GSB7 vs GSB6, indicating that the development of thorns underwent a lot of stimulus from late growth stage to browning stage (Supplemental Fig. S7). There were 134 pathways were significantly enriched (Q-value < 0.05) through KEGG pathway enrichment analysis (Fig. 5). The pathway ‘Plant hormone signal transduction’ were enriched in GSB2 vs GSB1 and GSB6 vs GSB5, elucidating that plant hormone played critical role in these four developmental stages (Fig. 5A and E). The pathway ‘Phenylpropanoid biosynthesis’ were enriched in GSB2 vs GSB1 and GSB5 vs GSB4, suggesting the significant role of phenylpropanoid in the thorns transition from formative stage to dormancy stage, and the stage of buds transforming to thorns to fast-growing stage (Fig. 5A and D). In GSB3 vs GSB2, DEGs were mainly enriched in ‘Biosynthesis of amino acids’ and ‘Ribosome’ pathways (Fig. 5B). In GSB4 vs GSB3, DEGs were mainly enriched in ‘Photosynthesis’ and ‘Carbon metabolism’ pathways (Fig. 5C). In GSB7 vs GSB6, DEGs were mainly enriched in ‘Starch and sucrose metabolism’ and ‘MAPK signaling’ pathways (Fig. 5F). These GO and KEGG enrichment analyses contributed to the understanding of regulatory pathways involved in the process of thorns development. Additionally, to validate the accuracy of RNA-seq data, the qRT-PCR analysis of randomly selected six DEGs confirmed that their expression trends were consistent with the RNA-seq, suggesting the reliablity of transcriptome data (Supplemental Fig. S8).

**Figure 5.**
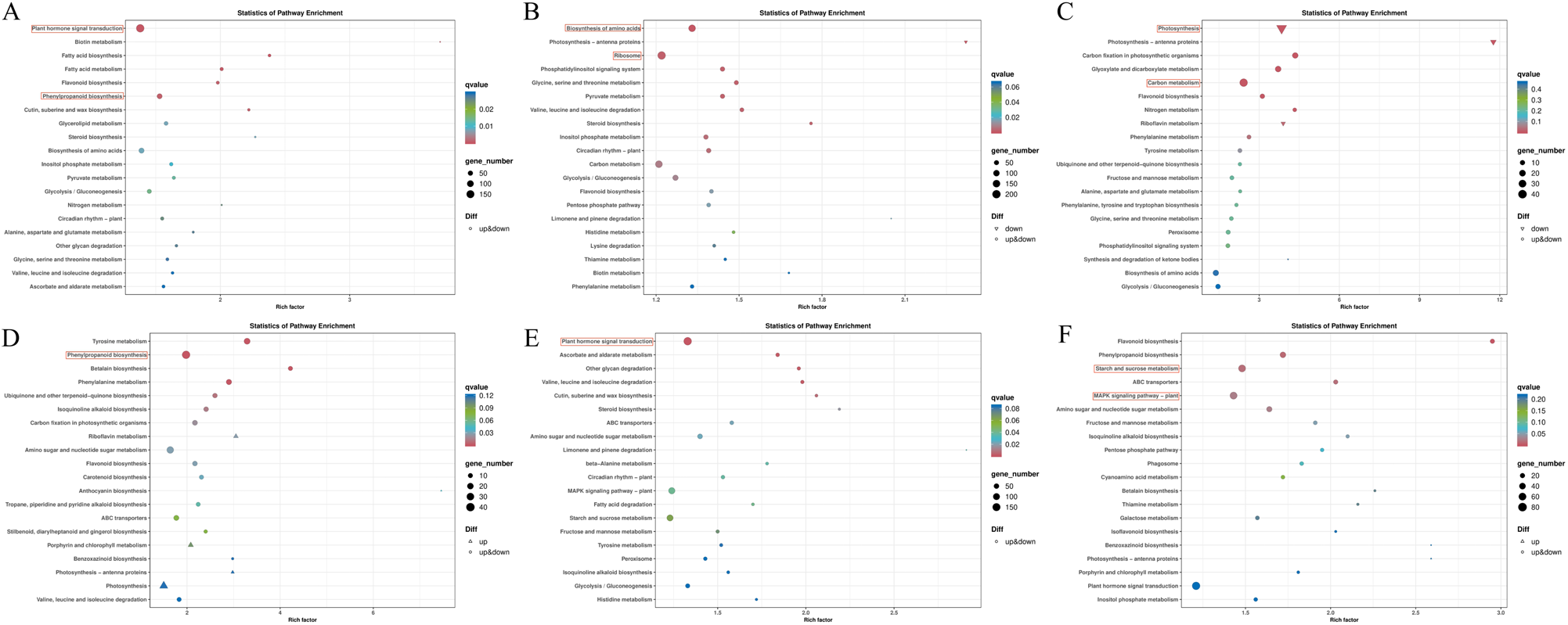
KEGG enrichment analysis in adjacent developmental stages. **A)** GSB2 vs GSB1, **B)** GSB3 vs GSB2, **C)** GSB4 vs GSB3, **D)** GSB5 vs GSB4, **E)** GSB6 vs GSB5, **F)** GSB7 vs GSB6.

### Hub genes involved in the thorn development in *G. sinensis*

To screen candidate transcripts and transcription factors (TFs) involved in the biosynthesis process of thorns in *G. sinensis*, the TFs were predicted among adjacent developmental stages. As a result, a total of eight genes encoding TFs were uncovered and classified into eight TF families, including *MYB*, *bZIP*, and *HB-HD-ZIP* (Supplemental Table S12). To further identify potential hub regulatory genes involved in thorns development, we performed weighted gene correlation network analysis (WGCNA) of DEGs with phenotype characters. Among the 13 identified modules, the expression pattern of honeydew1 module, including 137 DEGs, was tightly correlated with basal diameter (r = 0.51) and weight (r = 0.86). Additionally, the expression pattern of darkorange2 module, including 107 DEGs, was tightly correlated with length (r = 0.71) (Fig. 6, A and B). To further explore related key DEGs regulating thorns development, the gene regulation network was built using honeydew1 module and darkorange2 module. The GO annotation and KEGG enrichment analysis found that ‘Plant hormone signals’, ‘Plant pathogens’ and ‘Sucrose and starch metabolic’ pathways were more enriched in both two modules, indicating these pathways may be involved in regulating thorns development (Supplemental Fig. S9). Notably, the network presented that DEGs of honeydew1 module had high expression in GSB3-GSB5, including four transcription factors Gsin11G015360 (*MYB*), Gsin14G005090 (*HB-HD-ZIP*), Gsin01G010660 (*RLK-Pelle_LRR-XII-1*) and Gsin01G017400 (*RLK-Pelle_DLSV*) (Fig. 6, C and D). In addition, the DEGs of darkorange2 module had one transcription factors, including Gsin09G015570 (*bZIP*) and Gsin07G006200 (*RLK-Pelle_LRR-XI-1*) (Fig. 6, E and F). The above investigations suggested the hub roles of transcription factors such as *MYB*, *ZIP* should be involved in the thorns development of *G. sinensis*.

**Figure 6.**
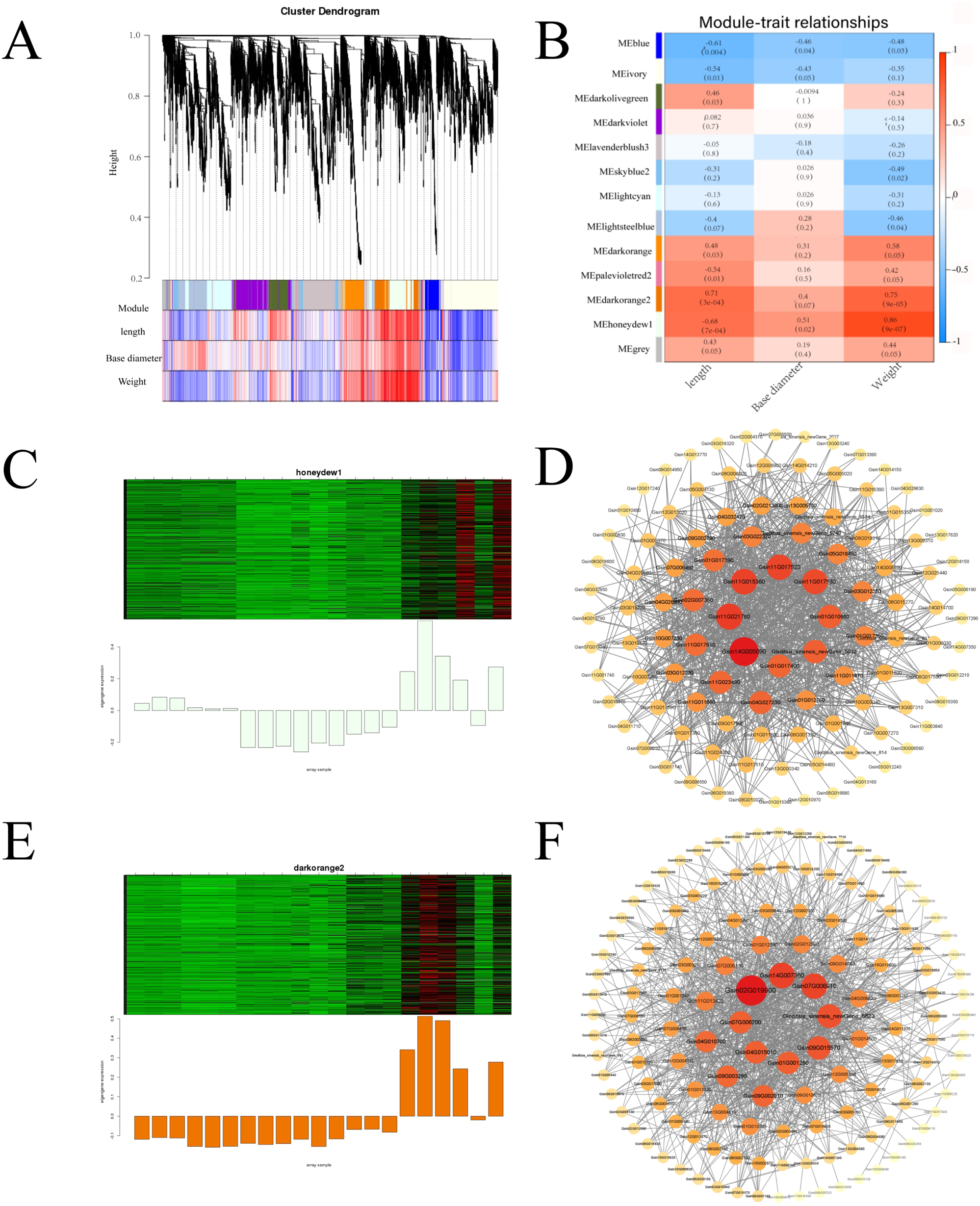
WGCNA analysis of genes related to the thorns development and gene regulatory network based on WGCNA in *Gleditsia sinensis*. **A)** Cluster dendrogram of identified modules. **B)** Module-trait relationship. **C)** and **D)** The network presents that DEGs of honeydew1 honeydew1 module. **E)** and **F)** The network presents that DEGs of darkorange2 module.

To investigate the potential roles of hub TFs involved in thorns development process, their expression profiling were identified. Gsin11G015360, Gsin14G005090, Gsin01G010660 and Gsin01G017400 were lowly expressed at GSB1-GSB5, while highly expressed at GSB5-GSB7 with a gradual rising trend, which indicated their potential regulation roles from fast-growing to browning development of thorns (Fig. 7A; Supplemental Fig. S10). Additionally, Gsin09G015570 and Gsin07G006200 were lowly expressed at GSB1-GSB5, while highly expressed at GSB5-GSB7, which indicated their potential regulation roles at the late growth stage of thorns (Fig. 7A; Supplemental Fig. S10).

**Figure 7.**
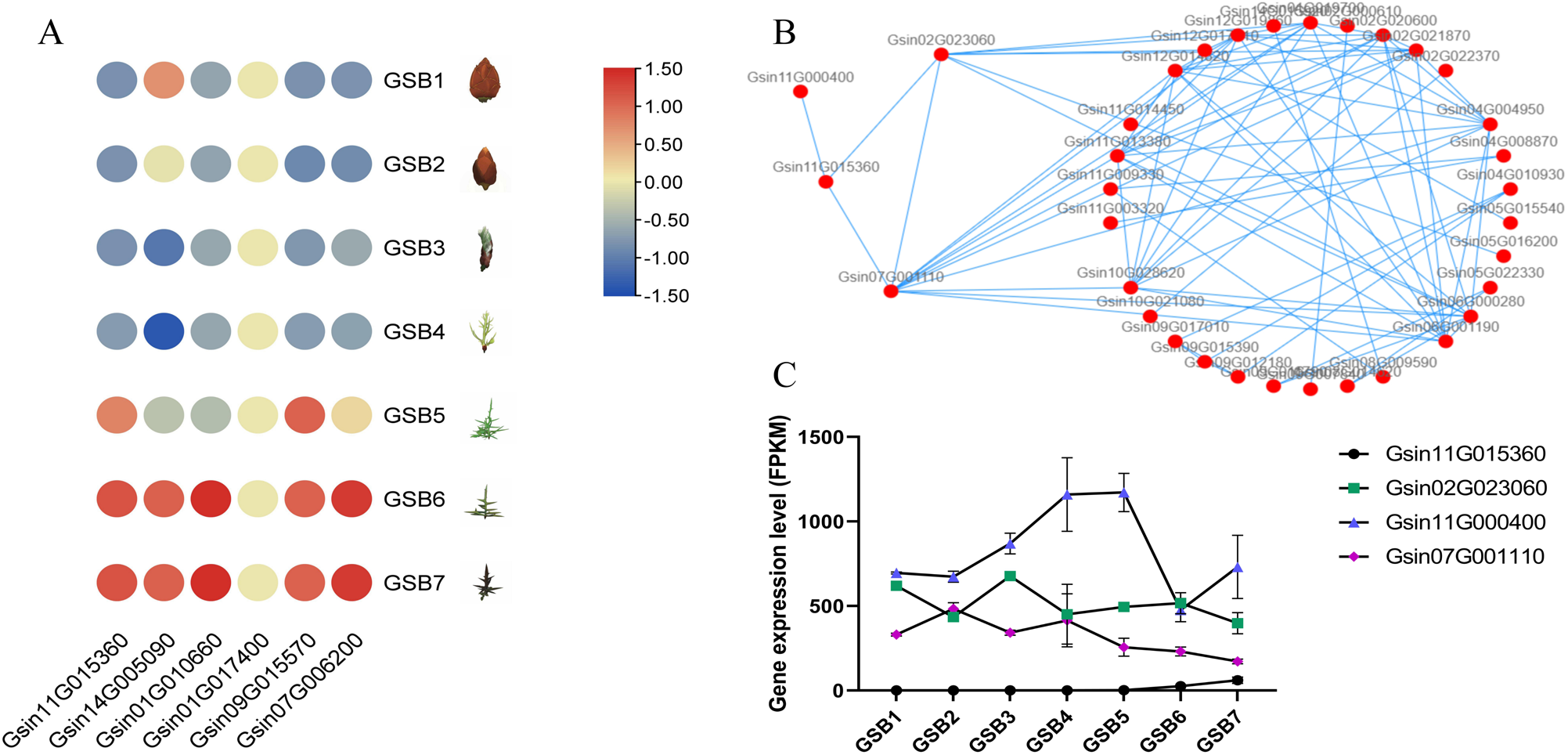
The expression patterns and interaction of hub genes involved in the thorns development in *Gleditsia sinensis*. **A)** The heatmap of 6 hub genes involved in thorns development at 7 developmental stages. **B)** The protein interaction prediction among Gsin11G015360 and its target genes (FPKM>100). **C)** The gene expression of Gsin11G015360 and its predicted protein interaction genes at 7 developmental stages.

### The potential molecular regulation mechanism of thorns development in *G. sinensis*

MYB had been widely reported in regulating the development of plants, including lateral root development, fruit ripening (Cao et al., 2020; Yang et al., 2020). To further analyze the potential molecular regulation mechanism of hub TFs *GsinMYB* (Gsin11G015360) in the thorns development of *G. sinensis*, its comparative highly expressed target genes (FPKM>100) were predicted. As a result, Gsin11G015360 might regulate the development of thorns by targeting 64 genes (FPKM>100). The GO enrichment analysis demonstrated that ‘Translation’ (GO:0006412), ‘Translational elongation’ (GO:0006414), ‘Glucose metabolic process’ (GO:0006006), ‘Nucleocytoplasmic transport’ (GO:0006913), ‘Photosynthesis’, ‘Light harvesting’ (GO:0009765), ‘Cell redox homeostasis’ (GO:0045454) and ‘Protein-chromophore linkage’ (GO:0018298) were significantly enriched (Supplemental Fig. S11A). KEGG pathway enrichment analysis showed that ‘Ribosome’ (ko03010), ‘Ubiquitin mediated proteolysis’ (ko04120), ‘Photosynthesis - antenna proteins’ (ko00196), ‘Arginine and proline metabolism’ (ko00330), ‘Carbon fixation in photosynthetic organisms’ (ko00710), and ‘Glycolysis / gluconeogenesis’ (ko00010) were significantly enriched (Supplemental Fig. S11B). Furthermore, the protein interactions among Gsin11G015360 and its target genes (FPKM>100) were predicted (Fig. 7B). It was found that Gsin11G015360 may interact with Gsin02G023060 (peptidylprolyl isomerase, Peptidyl-Prolyl cis / trans Isomerase (PPIase)), Gsin07G001110 (RNA-binding protein 8A), and Gsin11G000400 (high mobility group protein B3, HMG1 / 2-like protein isoform X1). The gene expression of Gsin02G023060 showed a trend of first rising and then falling. While the gene expression of Gsin07G001110 and Gsin11G000400 showed a gradual downward trend (Fig. 7C). Overall, Gsin11G015360 might be involved in the later development of thorns, and which might negatively interact with Gsin02G023060, Gsin11G000400, and Gsin07G001110. To further explore whether the Gsin11G015360 could regulate these interactive genes, the *cis*-acting regulatory elements (PLANTCARE; Supplemental Fig. S12) of 2000-bp promoter region in Gsin02G023060, Gsin07G001110, and Gsin11G000400 were analyzed. As a result, we identified many MYB TF binding sites in these three genes, which implied they may be the potential downstream targets of Gsin11G015360. Comprehensively, we speculate that the regulatory roles of Gsin11G015360 and its target genes in controlling the development of thorns, thereby contributing to the investigation of thorn breeding of *G. sinensis*.

## Discussion

Here we presented a high quality chromosome-level genome assembling of *G. sinensis* as Leguminosae species with important economic and medicinal value throughout the world. The full-length transcriptome sequencing of diverse tissues was performed to assist genome functional annotation. The comparative and evolutionary analysis of *G. sinensis* with eight Leguminosae species and two model plants unveiled the divergence of *G. sinensis* from other Leguminosae species. Through the integrated analyses of genome sequencing, phenotypic characteristics and gene expression profiling, we provided new insights into the genomic characteristics of *G. sinensis*, and its regulated genes involved in thorns development, identifying potentially useful hub genes for further studies of the genetic basis of molecular breeding of *G. sinensis*. Therefore, the *G. sinensis* genome assembly is of high reference for deciphering genome biology, deepening comparative genomic investigation and facilitating the genetic improvement of *G. sinensis* and related Leguminosae species.

### The divergence of *G. sinensis* from other Leguminosae species

The WGD event is widely present in plants and is an important event driving plant evolution. The increasing plant genomes have shown that they have experienced at least one WGD event. Genomic analysis of ancient polyploidization events showed almost all angiosperms were probably occurred around 66 million years ago (during the Cretaceous Period) (Zhang et al., 2020). Here, Ks and chromosome collinearity analysis revealed that all six Leguminosae species have underwent two WGD events, which was consistent with previous reports in the soybean genome (Schmutz et al., 2010). The first ancient polyploidization event of six Leguminosae species occurred approximately 58-66 million years ago (Zhang et al., 2020), which may be common to all core eudicots plants. In this investigation, the *G. sinensis* underwent WGDII event after differentiating from *M*. *truncatula* (Fig. 1D). The comprehensive analysis of phylogenetic tree and Ks suggested the first WGD event caused the differentiation of *G. sinensis* from other Leguminosae plants. Furthermore, the phylogenetic tree analysis revealed *G. sinensis* had the closest relationship with *A*. *hypogaea*, and they were differentiated approximately between 63.11 and 89.91 Ma. WGD is mainly reflected in the contraction and expansion of plant gene families and the generation of new endemic gene families (One thousand plant transcriptomes and the phylogenomics of green plants, 2019). Comparing to the 10 other plants, there were 559 unique gene families in *G. sinensis*, which may result in the production of thorns. The specific gene family enrichment analysis of *G. sinensis* found ‘Purine metabolism’ pathway, ‘RNA transport’ pathway and ‘Galactose metabolism’ pathway might play important roles in the formation and growth of thorns, which should promote the differentiation of *G. sinensis* from other Leguminosae species. The positive selection in genes is vital for the generation of new functions in species. We found 45 positive selection genes in the *G. sinensis* genome, which were mainly enriched in the ‘Protein export’ pathway. Among them, 23 genes were differentially expressed during seven developmental stages, suggesting they may play a regulatory role in the growth and development of thorns (Supplemental Fig. S13).

### Potential key pathways involved in thorns development

Light is crucial for the growth, morphology, and developmental changes in plants (de Wit et al., 2016). The photomorphogenesis was reported to participate in thorn development (Xiao et al., 2023). While the detailed role of light in regulating the development of thorns is largely unknown. The photosynthesis-antenna proteins have been verified to participate in the absorption and conversion of light energy. In this investigation, KEGG enrichment analysis in seven typical developmental stages found DEGs in ‘Photosynthesis-antenna proteins’ pathway were down-regulated at GSB3 vs GSB2 (Fig. 5B). Subsequently, the DEGs in ‘Photosynthesis’ and ‘Photosynthesis-antenna proteins’ pathway were down-regulated at GSB4 vs GSB3, while were up-regulated at GSB5 vs GSB4 (Fig. 5, C and D). These findings elucidated the critical role of light related pathways in early and mid-term development of thorns. The sucrose had been reported play a dual role of energy and signal in the occurrence of branch thorns (Lujia Li, 2023). Intriguingly, we found ‘Starch and sucrose metabolism’ pathway were enriched at both GSB6 vs GSB5 and GSB7 vs GSB6, suggesting its important role in the late development of thorns (Fig. 5, E and F). Plant hormone and phenypropanoid have been widely known involved in plant development (de Wit et al., 2016; Dong and Lin, 2021). Here, we found hormone related pathways were enriched at GSB2 vs GSB1 and GSB6 vs GSB5, and phenypropanoid related pathways were enriched at GSB2 vs GSB1 and GSB5 vs GSB4, indicating that plant hormone and phenypropanoid also regulating the thorns development (Fig. 5, A, D and E). Taken together, plant hormone, phenypropanoid, photosynthesis and saccharides were involved in thorn development in *G. sinensis* (Fig. 8).

**Figure 8.**
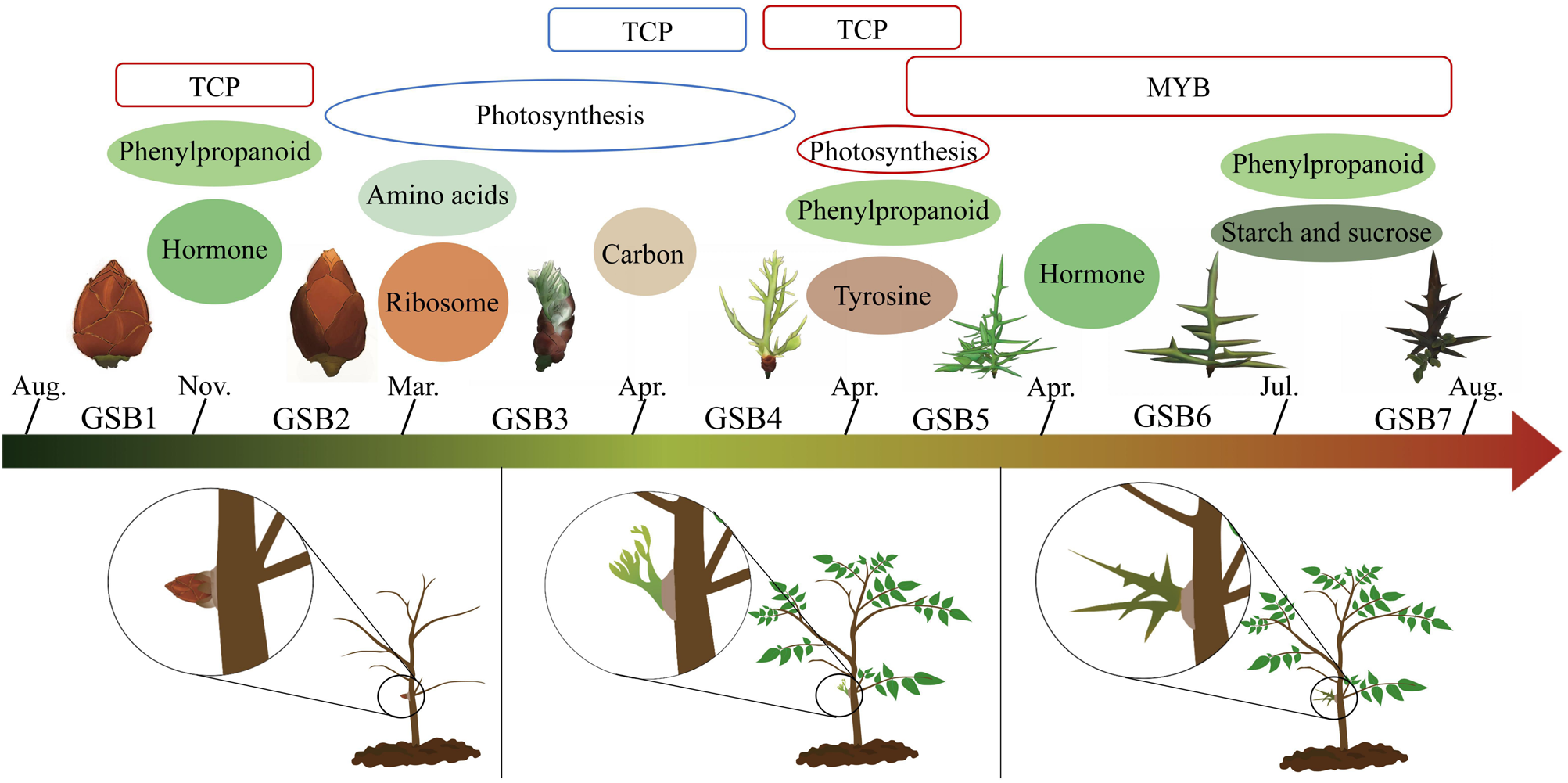
A proposed molecular regulation model of thorns development in *G. sinensis*. The blue outline represents a decrease in expression level, and red outline indicates an increase in expression level. Oval icons represent KEGG enrichment pathways in adjacent developmental stages. Roundrectangle icons represent transcription factors regulating the thorns development.

### Gene co-expression and potential TFs involved in thorns development

MYB TFs comprise one of the largest families of transcription factors in plants, which have been reported to play crucial roles in the evolution, development, and stress responses in plants (Zhu et al., 2019; Yang et al., 2022; Zhu et al., 2023). Here, the WGCNA analysis with phenotypic characters in this investigation found that *GsinMYB* (Gsin11G015360) might be involved in thorns development, especially the late developmental stage (Fig. 8). In this investigation, the expression of *GsinMYB*, the potential regulator involved in late growth of thorns, was the homologous gene of *ATMYB113*. Additionally, transcriptional activation activities of *ATMYB113* was found mediated the anthocyanin accumulation, the fruit color diferentiation (Bao et al., 2022; Li et al., 2023). Thus, we predicted that *GsinMYB* (Gsin11G015360) might regulate the color change from green to yellow brown and then to purple brown by mediating the anthocyanin accumulation. Similarly, earlier investigation proposed that *Myb-like*, *YABBY2*, *Growth-regulating factor 3*, *TCP2*, *Zinc transporter 8* may participate the maintenance and growth of thorns in *G. sinensis*, helping to elucidate MYB involved in thorns development (Xiao et al., 2023). In castor, the map-based cloning in 2 independent F_2_ populations presented the regulatory roles of *RcMYB106* about capsule spine development. Deeper analysis showed *RcMYB106* might target the downstream gene *RcWIN1* (*WAX INDUCER1*), and a series of experiments verified *RcMYB106* and *RcWIN1* might together act as positive regulators of capsule spine development in castor (Liu et al., 2023). Taken together, these investigations provide insights into the crucial role of MYB in the regulation of thorn development.

*TEOSINTE BRANCHED1 / CYCLOIDEA / PCF* (*TCP*), a plant-specifc transcription factor family, play central roles in many aspects of plant development, including plant height (Davière et al., 2014), lateral branching (Takeda et al., 2003), trichome formation (Lan et al., 2021), thorn conversion (Zhang et al., 2021) and circadian clock (Giraud et al., 2010). The gene expression profile analysis of *G. sinensis* seedlings showed TCP TFs might be involved in developmental initiation regulation of thorns (Xiao et al., 2023). Previous investigation elucidated two Citrus genes, *THORN IDENTITY1* (*TI1*) and *THORN IDENTITY2* (*TI2*), encoding TCP transcription factors, were critical regulators of the termination of meristem proliferation and concomitant thorn production (Zhang et al., 2021). The investigation in Citrus uncovered that the termination of *CsWUS* expression by *TI1* is critical for thorn identity, which further supported the vital roles of TCP TFs in thorn development. Consistently, we found 53 genes were noted as TCP TFs, among which Gsin09G001690 had the highest correlation with *TI1* / *2* in *Citrus* (Supplemental Fig. S14A). Furthermore, Gsin09G001690 exhibited a ‘M’ type expression trend during seven developmental stages, which meant Gsin09G001690 had the lowest expression level at GSB4 (the stage of buds transforming to thorns), suggesting the potential role of TCP in regulating the conversion of buds to thorns (Fig. 8; Supplemental Fig. S14B).

## Materials and methods

### Plant materials and sequencing

The terminal bud, young leaves, mature leaves, barks, pods and thorns of *G. sinensis* were collected from mature superior cultivars *Gleditsia sinensis* ‘Yulin No.1’ in September at Nanyang City, Henan Province, China (112.520270E, 33.507580N). The young leaves were used for genome sequencing and assembly. The genome size was estimated by second-generation sequencing. The whole genome was sequenced by Circular Consensus Sequencing (CCS) based on Sequel II platform, the long reads and high quality HiFi reads were used for subsequent investigation. The RNA-seq with Illumina platform supported gene prediction. The mix of samples were used for full-length transcriptome sequencing. The full-length cDNA of mRNA was synthesized using NEBNext Single Cell / Low Input cDNA Synthesis & Amplification Module. The full-length transcriptome sequencing was obtained with PacBio SMRT single molecule real-time sequencing technology based on sample mix.

The thorns of *G. sinensis* at formative stage (the latter part of October, GSB1), dormancy stage (the first part of January, GSB2), germination stage (the first part of April, GSB3), the stage of buds transforming to thorns (mid-April, GSB4), fast-growing stage (the latter part of April, GSB5), late growth stage (mid-June, GSB6) and browning stage (the first part of August, GSB7), were collected with three repeats in Beijing Forestry University (40.005875N, 116.347459E) for transcriptome sequencing. The samples were also restored in FAA fixative (50% ethanol : formaldehyde : glacial acetic acid : glycerol = 18 : 1 : 1 : 1) and electron microscope fixative respectively for observation of paraffin sections and electron microscopy. The genome and transcriptome sequencing procedures were performed by Bio Marker Technologies Company (Beijing, China).

### Genome assembly and assessment

The chromatin conformation capture (Hi-C) fragment libraries were constructed from 300-700 bp insert size and were measured by Illumina platform for auxiliary assembly (Rao et al., 2014). Hi-C data were filtered and evaluated by HiC-Pro (v2.10.0) (Servant et al., 2015). BWA (version: 0.7.17-r1188; align mode: aln; parameters: default) was used to align the sequencing double ended data with the sequence of the assembled genome. Subsequently, the ligating adjacent chromatin enables scaffolding in situ (LACHESIS) (paraments: CLUSTER_MIN_RE_SITES = 56; CLUSTER_MAX_LINK_DENSITY = 2; ORDER_MIN_N_RES_IN_TRUNK = 15; ORDER_MIN_N_RES_IN_SHREDS = 15) was used to obtain the version of chromosome level genome (Burton et al., 2013). Finally, the Benchmarking Universal Single-Copy Orthologs (BUSCO) v5.2.2 software was used to evaluate the accuracy and completeness of the assembled genome (Simão et al., 2015).

### Gene prediction and annotation

For repeat annotation, RepeatModeler2 (v2.0.1) (Flynn et al., 2020) was used to perform *de novo* sequencing based on RECON (v1.0.8) (Bao and Eddy, 2002) and RepeatScout (v1.0.6) (Price et al., 2005), and the prediction results were classified using RepeatClassifier (Flynn et al., 2020) using Dfam database (v3.5) (Wheeler et al., 2012); LTR_ Retriever (2.9.0) (Ou and Jiang, 2018) was used for the *de novo* prediction of LTR based on the prediction results of LTRharvest (v1.5.10) (Ellinghaus et al., 2008) and LTR_FINDER (v1.07) (Xu and Wang, 2007); the primary *de novo* repeat prediction results were combined with known database to obtain *G. sinensis* repeat sequences database, based on which, RepeatMasker (v4.1.2) (Tarailo Graovac and Chen, 2009) was used to predict transposable elements (TE). Additionally, tandem repeats were predicted by MIcroSAtellite identification tool (MISA v2.1) (Beier et al., 2017) and Tandem Repeat Finder (TRF, v409) (parameters: 2 7 7 80 10 50 500 -d -h) (Benson, 1999).

To predict the protein-coding genes, three strategies including *de novo*, homolog-based and transcriptome-based approaches were used. For *de novo* prediction, Augustus v3.1.0 (Stanke et al., 2008) and SNAP (Korf, 2004) were applied. For homolog-based prediction, the protein sequences of *A*. *thaliana*, *G*. *max*, *L*. *japonicus* and *M*. *truncatula* were aligned against the *G. sinensis* genome using GeMoMa v1.7 (Keilwagen et al., 2016). For transcriptome-based prediction, RNA-seq data were predicted by GeneMarkS-T v5.1 (Tang et al., 2015) PASA v2.4.1 (Haas, 2003). All the genes predicted by three methods were integrated and updated by EVM v1.1.1 (Haas et al., 2008) and PASA v2.4.1 respectively. The predicted protein-coding genes were functional annotated in databases including NR (ftp://ftp.ncbi.nlm.nih.gov/blast/db), EggNOG (Huerta-Cepas et al., 2019), GO (http://geneontology.org), KEGG (Kanehisa et al., 2016), SWISS-PROT (Boeckmann, 2003) and Pfam (Finn, 2006). The prediction of non-coding RNA were carried out through rRNA predicted in barrnap (v 0.9) (Loman, 2017), tRNA predicted in tRNAscan-SE (v1.3.1) (Lowe and Eddy, 1997), miRNA, snRNA and snoRNA predicted in Infenal 1.1 (Nawrocki and Eddy, 2013). The GenBlastA (version v1.0.4) was used to scan the whole genomes after masking predicted functional genes (She et al., 2009). Putative candidates were analyzed by searching for non-mature termination codon and frame-shift mutations using GeneWise (version 2.4.1) (Birney et al., 2004). The InterProScan (5.34-73.0) (Jones et al., 2014) was used for Motif annotation.

### Genome comparison and phylogenomics

The comparative genome analysis were performed among *Cicer arietinum* (ftp://ftp.ncbi.nlm.nih.gov/genomes/all/GCF/000/331/145/GCF_000331145.1_ASM33114v1), *Populus trichocarpa* (https://phytozome-next.jgi.doe.gov/info/Ptrichocarpa_v4_1), *Arachis hypogaea* (https://ftp.ncbi.nlm.nih.gov/genomes/all/GCF/003/086/295/GCF_003086295.2_arahy.Tifrunner.gnm1.KYV3/), *Lotus japonicus* (https://data.jgi.doe.gov/refine-download/phytozome?organism=Ljaponicus), *Phaseolus vulgaris* (https://phytozome.jgi.doe.gov/pz/portal.html#!bulk?org=Org_Pvulgaris), *Glycine max* (https://data.jgi.doe.gov/refine-download/phytozome?organism=Gmax), *Cajanus cajan* (https://ftp.ncbi.nlm.nih.gov/genomes/all/GCF/000/340/665/GCF_000340665.2_C.cajan_V1.1/), *Lupinus angustifolius* (https://ftp.ncbi.nlm.nih.gov/genomes/all/GCF/001/865/875/GCF_001865875.1_LupAngTanjil_v1.0/), *Arabidopsis thaliana* (https://phytozome-next.jgi.doe.gov/info/Athaliana_TAIR10), *Medicago truncatula* (https://ftp.ncbi.nlm.nih.gov/genomes/all/GCF/003/473/485/GCF_003473485.1_MtrunA17r5.0-ANR/) and *G. sinensis* genome. Orthofinder v2.4 software (Emms and Kelly, 2019) was applied to generate orthologous gene groups of gene families in 11 sequenced species, which was followed by annotation with PANTHER V15 database (Mi et al., 2019). The specific gene families in *G. sinensis* were analyzed by clusterProfile v3.4.4 (Yu et al., 2012) for GO and KEGG enrichment analysis. The phylogenetic tree was constructed with single copy gene sequence by IQ-TREE (v1.6.11) (Nguyen et al., 2015). The differentiation time was calculated with MCMCTREE program of PAML (v4.9i) (Yang, 1997). CAFE (v4.2) was used to analyze the expansion / contraction of gene families based on the chronogram of these 11 species and screening criteria that both family-wide *P*-Values and viterbi *P*-Values were lower than 0.05 (Han et al., 2013). CodeML program of PAML was used to investigate the positively selected genes among *G. sinensis*, *L*. *japonicus*, *C*. *cajan*, *G*. *max* and *L*. *angustifolius*.

### Genome synteny analysis

Diamond (v0.9.29.130) (Buchfink et al., 2015) and MCScanX (Wang et al., 2012) were used to identify syntenic blocks among *G. sinensis*, *L*. *japonicus*, *C*. *cajan*, *G*. *max* and *L*. *angustifolius*. The WGD events were identified with Ks based on wgd (v1.1.1) (Zwaenepoel and Van de Peer, 2019) and 4DTv (https://github.com/JinfengChen/Scripts). The long terminal repeat retrotransposons (LTR-RT) sequences were identified by LTR_FINDER (v1.07) (Xu and Wang, 2007) and LTRharvest (v1.5.9) (Ellinghaus et al., 2008). The LTR sequences were compared using MAFFT (parameters:--localpair --maxiterate 1000), and the distance K was calculated by Kimura model of EMBOSS (v6.6.0) (Rice et al., 2000).

### Phenotypic traits measurement and anatomical observations of *G. sinensis* thorns

The length, basal diameter and weight of seven thorns developmental stages were measured with a steel tape and electronic balance respectively. The thorns were fixed in FAA fixative for more than 24 h. Then fixed samples were dehydrated sequentially in an ethanol gradient series (75%, 85%, 90%, 95% and 100%), and treated with anhydrous ethanol II, alcohol benzene, then infiltrated with xylene I, xylene II, and embedded in paraffin. After solidification, the embedded sample was trimmed to trapezoid blocks. Sections of 4 μm without wax were stained with safranin and fast green and photographed with electric inverted fluorescence microscope (MDi8, Leica).

The samples of thorns were fixed in electron microscope fixative and osmic acid buffer for 2 h respectively. Subsequently, fixed samples were dehydrated with a gradual ethanol series (30%, 50%, 70%, 80%, 90%, 95% and 100%), isoamyl acetate dried by critical-point drying and coated with a gold layer, and observed with scanning electron microscope (SEM, SU8100, HITACHI).

### Transcriptome and analysis of differentially expressed genes among different developmental stages of *G. sinensis* thorns

Total RNAs of thorns at different development stages (GSB1-GSB7) were extracted for cDNA library construction. The cDNA libraries were sequenced on Illumina NovaSeq6000 sequencing platform. The filtered clean reads were mapped to the *G. sinensis* genome using HISAT2 (Kim et al., 2015). StringTie was applied to assemble the mapped reads, and reconstruct the transcriptome (Pertea et al., 2015). The novel transcripts and genes was achieved by StringTie and annotated by DIAMOND (Tatusov, 2000) against database including NR (Deng Y Y et al., 2006), Swiss-Prot (Apweiler, 2004), COG (Tatusov, 2000), KOG (Koonin et al., 2004), KEGG (Kanehisa, 2004). FPKM (Fragments Per Kilobase of transcript per Million fragments mapped) was applied to measure the expression level of a gene or transcript by StringTie using maximum flow algorithm (Trapnell et al., 2010). The differential expression analysis is performed by DESeq2, and define genes with absolute value of log_2_ fold change (FC) ≥ 2 and false discovery rate (FDR) < 0.01 as significant (Subramanian et al., 2005). GO, KEGG and WGCNA analysis was performed at BMK Cloud platform (www.biocloud.net).

### qRT-PCR verification of differentially expressed genes

Six DEGs were randomly selected for verification by qRT-PCR analysis with triplicates. The gene-specific primers were designed online (https://sg.idtdna.com/calc/analyzer). Real-time PCR amplifications were performed on the QuantStudio^TM^ Real-Time PCR Software using Hieff UNICON® Universal Blue qPCR SYBR Green Master Mix (YEASEN, 11184ES08). The relative expression level of genes was calculated using the 2^−ΔΔCt^ method and 18S rRNA was used as a reference gene.

### Accession numbers

Raw sequencing data was deposited in the China National GeneBank DataBase (CNGBdb) and can be accessed in the CNGB Sequence Archive. The accession number is CNP0005027.

## Acknowledgments

We thank Dr. Yingfeng Luo (BEIJING INSTITUTE OF GENOMICS CHINESE ACADEMY OF SCIENCES / CHINA NATIONAL CENTER FOR BIOINFORMATION) for the advice on this manuscript. No conflict of interest is declared.

## Author contributions

Y. L., D. F., F. L. and Y. W. designed this study; D. X. wrote the manuscript; J. L. performed the genome database investigation, J. W. performed the RNA-seq database investigation, Y. Y., Y. L. and J. W. collected plant materials, X. Y. revised the manuscript, R. Y. and C. W. performed the data curation. All authors have read and approved the manuscript.

## Supplemental data

The following materials are available in the online version of this article.

**Supplemental Figure S1.** K-mer based estimation of genome characters of *Gleditsia sinensis*.

**Supplemental Figure S2.** Genome-wide chromatin interaction heatmap of *Gleditsia sinensis* based on Hi-C data. The horizontal and vertical coordinates represent the Order of each bin on the corresponding chromosome group respectively.

**Supplemental Figure S3**. GO A) and KEGG B) enrichment analysis of expanded gene families in *Gleditsia sinensis*.

**Supplemental Figure S4.** Chromosome collinearity analysis between *Gleditsia sinensis* and *Arachis hypogaea* **A)**, and between *Gleditsia sinensis* and *Glycine max* **B)**, *Lupinus angustifolius* **C**), *Lotus japonicus* **D)**, and *Medicago truncatula* **E)**.

**Supplemental Figure S5.** The chart of gene expression of *Gleditsia sinensis* thorns at different stages. **A)** Comparison of FPKM density distribution, **B)** FPKM boxplot, **C)** Heat map of correlation analysis, **D)** PCA analysis.

**Supplemental Figure S6.** The analysis of differentially expressed genes between different stages in *Gleditsia sinensis* thorns. G1: GSB2 vs GSB1; G0: GSB3 vs GSB2; G2: GSB4 vs GSB3; G3: GSB5 vs GSB4; G4: GSB6 vs GSB5; G5: GSB7 vs GSB6.

**Supplemental Figure S7.** GO annotation analysis at adjacent developmental stages. **A)** GSB2 vs GSB1, **B)** GSB3 vs GSB2; **C)** GSB4 vs GSB3; **D)** GSB5 vs GSB4; **E)** GSB6 vs GSB5; **F)** GSB7 vs GSB6.

**Supplemental Figure S8.** The qRT-PCR analysis of randomly selected six differentially expressed genes.

**Supplemental Figure S9.** The KEGG pathway analysis of honeydew1 and darkorange modules.

**Supplemental Figure S10.** The genes expressed level of key transcription factors in different development stage of *Gleditsia sinensis* thorns.

**Supplemental Figure S11.** GO **A)** and KEGG **B)** enrichment analysis of target genes (FPKM>100) of hub gene Gsin11G015360.

**Supplemental Figure S12.** The *cis*-acting regulatory elements of predicted protein interaction genes of Gsin11G015360. **A)** The *cis*-acting regulatory elements of Gsin02G023060. **B)** The *cis*-acting regulatory elements of Gsin07G001110. **C)** The *cis*-acting regulatory elements of Gsin11G000400.

**Supplemental Figure S13.** The KEGG enrichment analysis of positively selected genes in *Gleditsia sinensis*.

**Supplemental Figure S14.** The analysis of transcription factor TCP in *Gleditsia sinensis.* **A)** The heatmap of the correlation between predicted TCP transcription factors in *Gleditsia sinensis* and TI1 / 2 in *Citrus*. **B)** The expressed level of TCP in different development stage in *Gleditsia sinensis* thorns.

**Supplemental Table S1.** Hi-C assisted assembly of *Gleditsia sinensis* 14 chromosome data.

**Supplemental Table S2.** Busco results for *Gleditsia sinensis* genome.

**Supplemental Table S3.** TE sequence prediction statistical results in *Gleditsia sinensis* genome.

**Supplemental Table S4.** Statistical results of tandem repeats in *Gleditsia sinensis* genome.

**Supplemental Table S5.** Statistical results of genes in *Gleditsia sinensis* genome.

**Supplemental Table S6**. Statistical results of non-coding RNAs in *Gleditsia sinensis* genome.

**Supplemental Table S7.** Prediction of pseudogenes in *Gleditsia sinensis* genome.

**Supplemental Table S8.** Statistics of gene function annotation statistics in *Gleditsia sinensis* genome.

**Supplemental Table S9.** Statistical analysis of homologous gene families in 11 sequenced species.

**Supplemental Table S10.** Chromosome collinearity analysis of intra-genome and inter-genome.

**Supplemental Table S11.** RNA-Seq data statistics.

**Supplemental Table S12.** Prediction of transcription factors in adjacent developmental stages of *Gleditsia sinensis*.

